# HAT: *de novo* variant calling for highly accurate short-read and long-read sequencing data

**DOI:** 10.1101/2023.01.27.525940

**Authors:** Jeffrey K. Ng, Tychele N. Turner

## Abstract

**Motivation:** *de novo* variant (DNV) calling is challenging from parent-child sequenced trio data. We developed **H**are **A**nd **T**ortoise (HAT) to work as an automated workflow to detect DNVs in highly accurate short-read and long-read sequencing data. Reliable detection of DNVs is important for human genetics studies (e.g., autism, epilepsy).

**Results:** HAT is a workflow to detect DNVs from short-read and long read sequencing data. This workflow begins with aligned read data (i.e., CRAM or BAM) from a parent-child sequenced trio and outputs DNVs. HAT detects high-quality DNVs from short-read whole-exome sequencing, short-read wholegenome sequencing, and highly accurate long-read sequencing data.

**Availability:** https://github.com/TNTurnerLab/HAT

**Supplementary information:** Supplementary data are available at bioRxiv.

## 1 Introduction

*de novo* variants (DNVs) are variants present in children but not in their parents. On average, humans have ~40-100 DNVs within their genome. Characteristics of DNVs genome-wide include approximately 20% occur at CpG sites and approximately 75% arise on the paternal chromosome of origin (Ng, et al., 2022). To date, DNV calling and methods for DNV calling have primarily focused on whole-exome sequencing (WES) and whole-genome sequencing (WGS) from short-read technologies (i.e., Illumina) (Allen, et al., 2013; DDD, 2017; Iossifov, et al., 2014; Turner, et al., 2017; Werling, et al., 2018). This assessment has enabled reliable detection of DNVs from regions of the genome with good mappability (i.e., unique regions). Newer highly accurate long-read sequencing data (i.e., Pacific Biosciences HiFi) is providing novel insights into the more challenging repeat regions of the genome (Wenger, et al., 2019). In this paper, we introduce Hare-And-Tortoise (HAT) as a *de novo* variant caller for sequencing data from short-read WES, short-read WGS, and long-read WGS in parent-child sequenced trios. HAT is important for generating DNV calls for use in studies of mutation rates (Ségurel, et al., 2014) and identification of disease-relevant DNVs (Iossifov, et al., 2014). Unlike the majority of DNV callers, the ability to call DNVs from multiple sequencing types is unique.

## 2 Application

### Datasets

Short-read WES data for 100 trios from the SPARK collection (SPARK, 2018) were assessed using HAT. Short-read WGS data were assessed from our previous publication in 4,216 trios from the Simons Simplex Collection (Ng, et al., 2022). Highly accurate long-read WGS data for four trios were assessed from our previous publications (Mehinovic, et al., 2022; Sams, et al., 2022).

The general HAT workflow consists of three main steps: GVCF generation, family-level genotyping, and filtering of variants to get final DNVs (Supplemental Figure 1). We leveraged the power of the GPU accelerated NVIDIA Parabricks (Franke and Crowgey, 2020) software (v4.0.0-1) https://docs.nvidia.com/clara/parabricks/4.0.0/index.html for rapid GVCF generation in Hare. This version of Parabricks does not require the purchase of a license, making it free to use for all with a GPU equipped system that is able to access and run Docker (Merkel, 2014) images. We specifically use the Parabricks versions of GATK HaplotypeCaller (McKenna, et al., 2010; Poplin, et al., 2018) and DeepVariant (Poplin, et al., 2018) for GVCF generation. We also provide a purely CPU based option, Tortoise, which uses the open-source versions of both programs. The genotyping step is done with GLnexus (Yun, et al., 2020) and post-genotyping filtering is done with our custom workflow.

HAT is capable of running on docker compatible machines and high-performance clusters (HPCs), as well as in the cloud. We offer the workflow as a Snakemake (Koster and Rahmann, 2012) workflow and a Cromwell workflow https://cromwell.readthedocs.io/en/stable/. Four V100 or A100 GPUs are recommended to run Parabricks on WGS data, with an average run time of ~40 minutes on our HPC. The total run time for Hare with WGS data, assuming four GPUs, is 4.5 hours. This runtime can be sped up with mass parallelization. We also ran Hare in the cloud using the Google Cloud Platform and the general cost for a 30x WGS trio was $8.54 USD with an overall runtime of about four hours.

Specific features of HAT to run on WES data include a specific post filtering script that separates the results into high and low confidence regions (Supplemental Figure 2). By default, it trims the capture region by 40 bp on both sides, however, this value can be adjusted to fit varying capture regions. Only one GPU is needed to accelerate GVCF generation to about four minutes, with a total parallelized run time of seven minutes.

Specific features of HAT to run on PB-WGS data include an optimized Tortoise to run on PacBio data with the ability to switch the model type for DeepVariant “PACBIO.” The overall runtime is 2.5 days.

## 3 Results

### DNVs from short-read WES

We tested HAT on 100 trios from the SPARK collection. After manual confirmation by visual inspection of the underlying read data, we saw that many of the false positive DNVs were found to be near the beginning or ends of the capture region. To address this, we added another filtering step that trims the provided capture region by 40 bp on each end, by default, and then sorts the DNVs into what we define as high and low confidence calls (Supplemental Figure 2). Assuming the capture region has a buffer of 50 bp on each end, we consider DNVs found within the capture region ± 10 bp to be in the high confidence category. DNVs found within the 40 bp sections at each end of the capture region are in the low confidence category. Within the high confidence category, we saw a DNV confirmation rate of 91.6%, as compared to 70.7% in the low confidence category. Features of high confidence category DNVs are 2.16 ± 1.29 DNVs per individual, 34.7% within CpG regions, and a Ti/Tv ratio of 2.39. We find all of these metrics to be within our expectations when calling DNVs from WES data.

### DNVs from short-read WGS

We previously tested HAT on 4,216 trios, with DNA derived from blood, from the Simons Simplex Collection (SSC) (Ng, et al., 2022). Overall, we identified a total of 329,589 DNVs with 78 ± 15 DNVs per individual. We found 18 ± 4.7% of the DNVs appear in CpG sites. Lastly, we observed a Ti/Tv ratio of 2.11. All of these values fall in line with our expectations. Overall, we showed that this pipeline is highly capable of detecting DNVs from short-read WGS.

### DNVs from long-read WGS

We ran HAT on four different long-read sequenced trios (Figure 1A & 1B) to test the advocacy of our pipeline on this newer sequencing technology. After filtering, we found ~94 per trio within unique regions of the genome (Supplemental Table 1). Long-read sequencing allows for more accurate DNV detection in repeat regions (Noyes, et al., 2022). Therefore, we tested whether HAT could reliably identify DNVs within repeats. HAT was able to detect ~62 per individual from these regions, with a total of ~156 DNVs found per individual (Supplemental Table 1). After manual inspection of the underlying read data for each of these DNVs, we saw a confirmation rate of 87.3% in the unique regions of the genome, 69.3% in the repeat regions, and an overall confirmation rate of 80.5% (Supplemental Table 2). This was much lower than our initial expectations. When assessing fold coverage of the genome in the families, the 9p.100 family had the highest confirmation rate as well as being the most deeply sequenced, with the parents’ coverage ~32x and the child at 46.1x, compared to the ~20x coverage of the PB.100 family (Figure 1A and Figure 1B). This family had DNV metrics within our expectation for long-sequencing reads, with 125 DNVs detected across the genome, 21.1% were found in CpG regions, and a Ti/Tv ratio of 2.11. From this analysis, we hypothesized our lower-than-expected DNV confirmation rate was due to the lower coverage seen in the PB.100 family in comparison to 9p.100.

**Figure 1).**
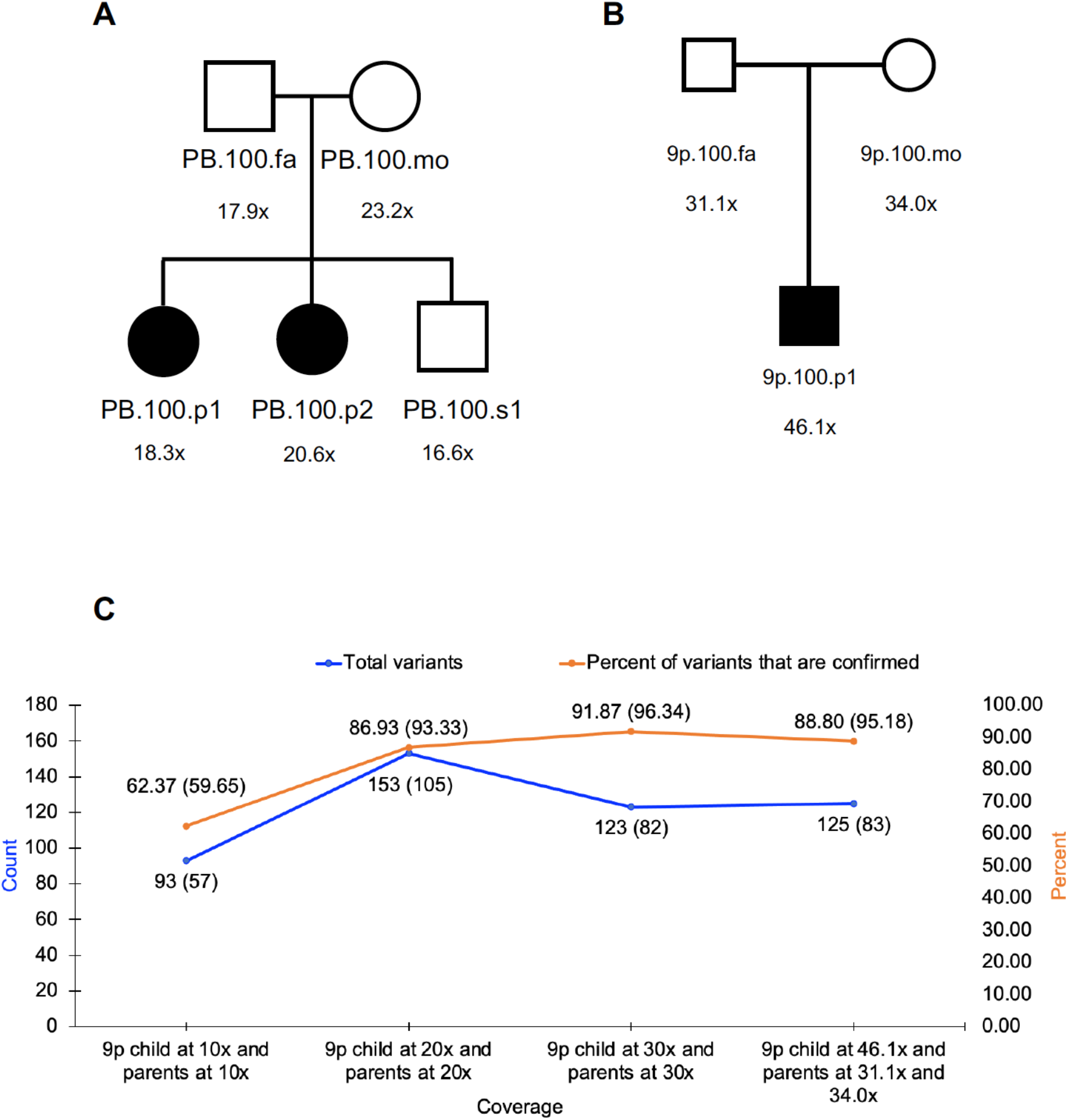
Long-read sequencing families and DNVs called with HAT. A) Pedigree of the PB.100 family, with long-read sequencing coverage shown. This family was sequenced by Pac-Bio HiFi sequencing in Mehinovic et al. 2022. B) Pedigree of the 9p. 100 family, with long-read sequencing coverage shown. This family was sequenced by PacBio HiFi sequencing in Sams et al. 2022. C) This graph illustrates the increase in the quality of DNV calls as the coverage increases in the downsampled 9p.100 family. The percent of DNVs confirmed, shown in orange, increases with coverage. The percentage in parenthesis is the percent of confirmed DNVs in unique regions of the genome. The total DNV count, shown in blue, is around expected as coverage increases. The counts in parenthesis are the confirmed DNVs in unique regions of the genome.

To test this hypothesis, we downsampled the 9p. 100 family to ~10x, ~20x, and ~30x for each member and ran HAT on these samples. As the coverage increased from 10x to 30x, the confirmation rate increased from 62.4% to 91.9% in 30x coverage (Figure 1C) with a confirmation rate of 96.3% in variants residing in unique regions of the genome. From these experiments, we conclude that 30x coverage genomes, in each of the members of the parent-child sequenced trio, are required for accurate DNV calling from highly accurate long-read WGS data.

## 4 Discussion

DNV calling from multiple sequencing types is critical for studies of mutation rates and human disease. Currently, the majority of sequencing data utilized for assessing DNVs is from short-reads. However, we are at a juncture in genomics whereby highly accurate long-read sequencing data will become more commonplace and a method to assess DNVs in this type of data is especially critical. We show that our method, HAT, works on both short-read and long-read data by advancing it to work on short-read WES and long-read WGS datasets in addition to short-read WGS. This workflow will be of interest to individuals studying mutation rates and human disease, respectively.

## Acknowledgements

Thank you to members of the Turner Laboratory at Washington University in St. Louis for helpful discussions on this work. We thank Elvisa Mehinovic for her help in manual inspection of the underlying read data in the 100 WES families.

## Data Availability

The main HAT GitHub repo can be found here: https://github.com/TNTurnerLab/HAT. The SSC WGS data are available as described previously (Ng, et al., 2022). The PB.100 and 9p.100 family datasets are available as described previously (Mehinovic, et al., 2022; Sams, et al., 2022).

