## Supplemental for "HAT: *de novo* variant calling for highly accurate short-read and long-read sequencing data"

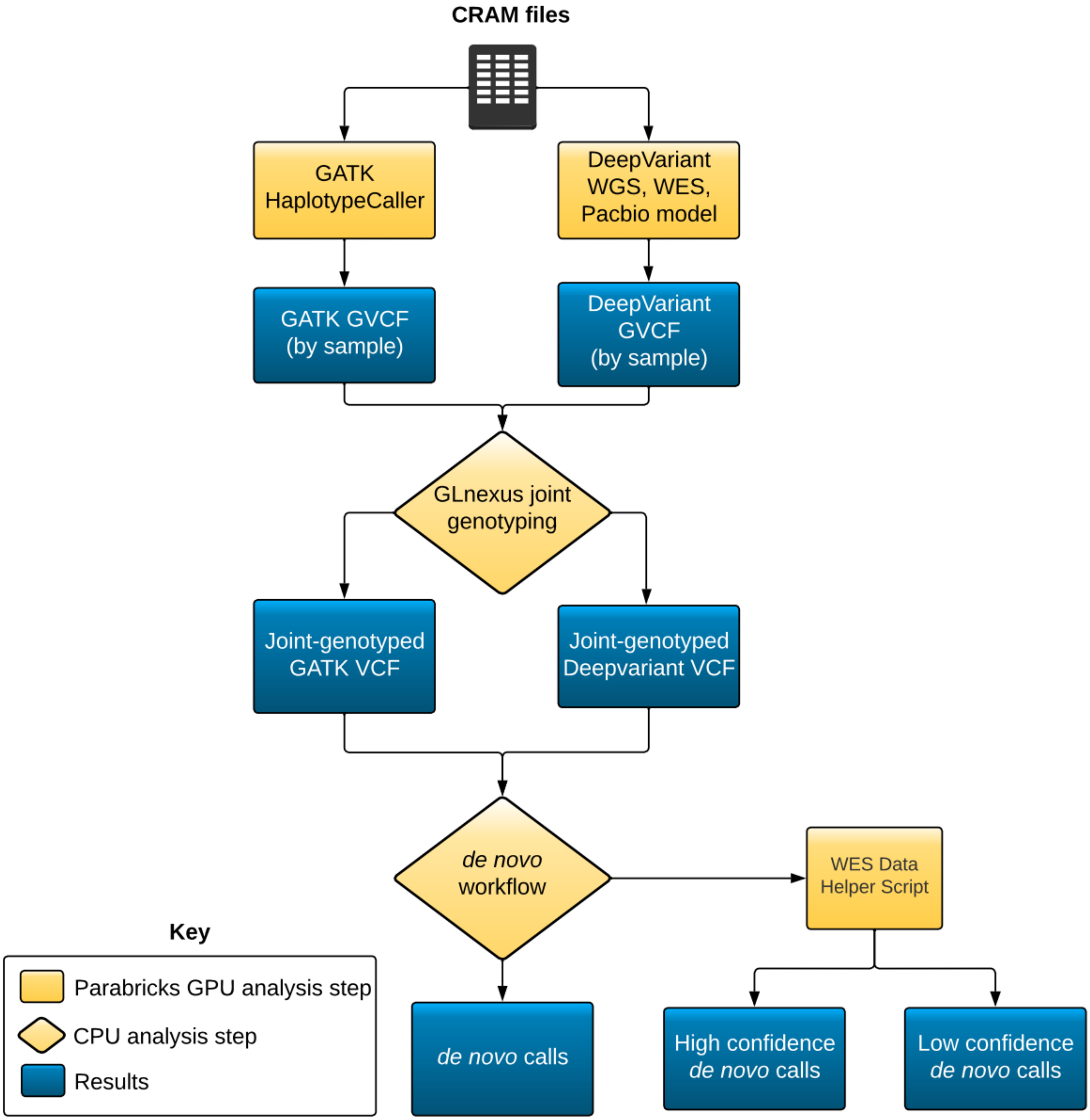


**Supplemental Figure 1 HAT workflow schematic.**
This figure shows how the general workflow of how HAT works.

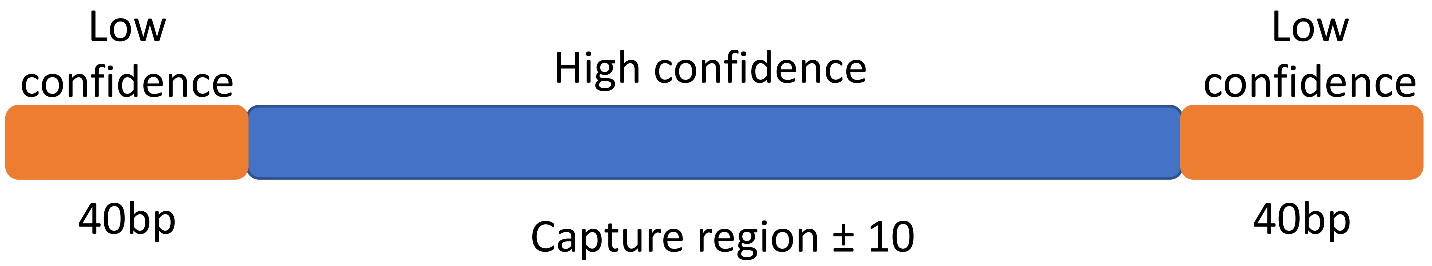


**Supplemental Figure 2 Filtering of WES capture regions and definition of high and low confidence DNV regions.**
This figure shows how we partition the provided capture regions to define high and low confidence DNVs. Any DNVs found within the capture region, plus 10bp on each side, would be considered high confidence DNVs and any DNVs in the 40bp at each end would be low confidence.

| Sample | Total *de novo* variants | *de novo* variants in unique regions of the genome | *de novo* variants in repeats |
|---|---|---|---|
| PB.100.p1 | 191 | 115 | 76 |
| PB.100.p2 | 174 | 87 | 87 |
| PB.100.s1 | 134 | 92 | 42 |
| 9p.100.p1 | 125 | 83 | 42 |

**Supplemental Table 1 *de novo* variants detected in four families sequenced with long-read sequencing.** This table shows the number of detected *de novo* variants (DNVs) for four long-read sequenced families. The blue indicates a greater amount of variants, red lower.

| Sample | Percent confirmed total *de novo* variants | Percent confirmed *de novo* variants in unique regions of the genome | Percent confirmed *de novo* variants in repeats |
|---|---|---|---|
| PB.100.p1 | 83.25 | 89.57 | 73.68 |
| PB.100.p2 | 77.01 | 83.91 | 70.11 |
| PB.100.s1 | 73.13 | 80.43 | 57.14 |
| 9p.100.p1 | 88.80 | 95.18 | 76.19 |

**Supplemental Table 2 Percent of confirmed *de novo* variants detected in four families sequenced with long-read sequencing.**
This table shows the percent of confirmed DNVs for the four long-read sequenced families. The blue indicates a high percentage, red lower. DNVs were manually verified via visual inspection of the reads.
